# Disrupted Cholesterol Biosynthesis and Hair Follicle Stem Cell Impairment in the Onset of Alopecia

**DOI:** 10.1101/2023.11.07.566136

**Authors:** Leemon Nikhila, Suresh Surya, Shahul Hameed Najeeb, Thankachan Mangalathettu Binumon, Aayush Gupta, Sandeep Gopalakrishnan, Parameswara Panicker Sreejith

## Abstract

**Background:** Hair follicle cycle and the functioning of stem cells in alopecia are influenced by the suppression of cholesterol synthesis and the accumulation of sterol intermediates.

**Objective:** This investigation aims to elucidate the regulatory function of disrupted cholesterol homeostasis in the functioning of hair follicle stem cells (HFSCs) and the cycling of hair follicles. Additionally, it seeks to provide an understanding of the fate of stem cells in primary cicatricial alopecia (PCA).

**Methods:** To evaluate the influence of cholesterol on the functionality of hair follicles, a study was conducted to analyse gene expression and pathways associated with hair follicle stem cell markers in scalp samples affected by PCA (LPP, FFA, CCCA, DC, DF, TF). To assess the influence of disrupted cholesterol homeostasis on HFSCs, we conducted experiments involving the administration of 7-dehydrocholesterol (7DHC) and BM15766 (Pharmacological inhibitor of cholesterol biosynthesis), to Human Hair Follicle Outer Root Sheath Cells (HORSCs), as well as C57BL/6 mice, and hair follicle organoid cultures. The study utilised reverse transcription polymerase chain reaction (RT-PCR) to assess gene expression, while immunofluorescence was employed to analyse protein expression. The tracking of stem cell fate was accomplished through the utilisation of a BrdU pulse-chase experiment, while the verification of apoptotic consequences was established by utilising the TUNEL assay. A statistical analysis was conducted to assess the statistical significance of the data.

**Results:** There was a notable decrease in the expression of HFSC marker genes among patients afflicted with PCA. *In vitro* data further confirmed the cholesterol inhibition and sterol intermediate accumulation in stem cells, leading to stem cell characteristics’ disruption. The experimental group of mice exposed to 7DHC and BM15766 also exhibited a notable inability to initiate hair regrowth. Consequently, this deficiency in hair regrowth resulted in the activation of apoptosis, specifically in the stem cells. Additionally, our confirmatory analysis, which was performed utilising organoid culture, consistently yielded comparable results. The results as mentioned above emphasise the significant importance of cholesterol production in preserving the integrity and functionality of HFSCs, hence providing novel insights into the progression of alopecia.

**Conclusion:** Individuals with aberrant cholesterol production, particularly those afflicted with PCA, experience a persistent impairment in hair regrowth due to the irreversible destruction of their hair follicles. The observed phenomenon is hypothesised to be attributed to the loss of hair follicle stem cells. Our study presents additional findings that elucidate the previously unknown involvement of sterol intermediates in regulating hair follicle cycling and stem cell function in PCA. The regulation of cholesterol production and the buildup of sterol intermediates have an impact on the hair follicle cycle and the role of stem cells in alopecia.

## 1. Introduction

The skin is a complex organ with appendages like hair follicles, sweat glands, sebaceous glands, and nails (Lee *et al.,* 2018). Stem cells are crucial in regenerating these appendages and supporting skin health. Hair follicles, among these, are essential for skin balance and have significant psychological importance. Hair disorders can affect growth, structure, and appearance, impacting self-esteem and quality of life (Roh & Lyle.,2006). Understanding the mechanisms behind these disorders is crucial for accurate diagnosis and effective treatment, enhancing patient outcomes.

Alopecia is a persistent dermatological condition characterized by progressive hair loss from the scalp and other body areas. It is an inflammatory disorder primarily affecting hair follicles. The aetiology and pathogenesis of this condition remain incompletely understood; however, it is widely acknowledged as an autoimmune disorder influenced by hereditary and environmental factors (Madani & Shapiro 2000). Primary cicatricial alopecia (PCA) is presently managed as an inflammatory condition that results in the irreversible damage of hair follicles. Nevertheless, the therapy options currently accessible for PCA are somewhat restricted and frequently prove inadequate, resulting in a slowdown in the disease’s advancement. (Panicker *et al.,* 2012). Primary cicatricial alopecia is classified into three categories based on the predominant inflammatory cell detected during the active phase: lymphocytic (e.g., Lichen Planopilaris, Frontal Fibrosing Alopecia, Central Centrifugal Cicatricial Alopecia, pseudopelade of Brocq), neutrophilic (e.g., Folliculitis Decalvans, Tufted Folliculitis), and mixed (e.g., Dissecting Cellulitis, acne keloidalis nuchae) (Mirmirani *et al.,* 2005).

Though the exact cause remains elusive, inflammation and the arrival of immune cells are distinctive features of PCA. Recent findings have hinted at the potential involvement of disrupted lipid balance in this context. Connections have been established between changes in cholesterol levels and PCAs (Palmer *et al.,* 2020). When 7-DHCR is inhibited or exogenous 7-dehydrocholesterol (7-DHC) is introduced to human primary outer root sheath keratinocytes or applied topically to mouse back skin, it produces a pro-inflammatory response. This reaction involves the enhancement of Toll-like receptor and interferon signalling pathways. Additionally, inhibiting cholesterol biosynthesis leads to an elevated TGFβ (Panicker *et al*., 2012), a widely recognized trigger of catagen (hair follicle regression) (Hibino & Nishiyama., 2004) and fibrosis (scarring) (Imanishi *et al.,* 2018).

Hair shaft regeneration from hair follicles (HFs) hinges on activating a crucial pool of hair follicle stem cells (HFSCs). This essential mechanism is central to the use of stem cell-based treatment for alopecia. Hair follicle stem cells are located in specific regions within the hair follicle structures like the bulge region (Cotsarelis *et al.,* 1990), lower permanent portion (hair germ) (Li *et al.,* 2003), and upper portion of the follicle (infundibulam) (Jahoda & Renolds., 2001). The studies suggest that the bulge area of the follicle is an anatomical niche for the stem cells and is capable of self-renewal (Waters *et al.,* 2007). The irreversible damage to the hair follicle stem cell destroys the regenerative capacity of the hair follicle (Harries *&* Paus., 2009). In studies conducted in transgenic mice, the targeted deletion of epithelial hair follicle stem cells results in the disappearance of the hair follicles (Ito *et al.,* 2005). Hair follicle stem cell regeneration depends on a complex network orchestrated by multiple pathways, including the Wnt/β-catenin pathway, Sonic hedgehog (Shh) pathway, Notch pathway, BMP (bone morphogenetic proteins) pathway, and apoptotic pathway. Among these, the Wnt/β-catenin signaling pathway emerges as a pivotal player, with its diverse functions in initiating hair follicle stem cells (HFSCs) activation for promoting hair growth during hair regeneration (Cleavers et al., 2014). Cholesterol modifications are also vital for signal transduction in the Wnt-β-catenin and hedgehog pathways (Incardona and Eaton 2000), both of which are fundamental in the control of human hair follicle (HF) cycling (Lee and Tumbar 2012). Therefore, cholesterol is crucial for normal hair follicle biogenesis and enhancing stem cell regeneration. A deeper comprehension of the disrupted cholesterol homeostasis within the realm of HFSC function can unlock new solutions for individuals grappling with persistent hair loss due to PCA. Our quest to shed light on the substantial role of the cholesterol biosynthesis pathway in HFSC function represents a significant stride towards revitalizing hair follicles in those affected by alopecia.

Additionally, our research identifies that modifications in the synthesis of cholesterol in the cells of hair follicles can initiate apoptosis of HFSC, resulting in the irreversible deterioration of hair follicles in mouse skin models and individuals with PCA. The findings mentioned above elucidate a hitherto unrecognised function of cholesterol precursors in the modulation of HFSC activity and their involvement in the pathogenesis of PCA. Additionally, this study demonstrates a novel association between sterols and the functioning of stem cells, which has the potential to significantly transform the field of diagnosing and treating these conditions.

## 2. Materials and methods

### 2.1. Human tissue

All human subjects involved in this study received explicit approval from the Human Research Ethics Committee of the University of Kerala (Protocol Number: ULECRIHS/UOK/2018/35). Written informed consent was secured following ethical protocols for research involving human participants. The identification of PCA was established through the examination of clinical observations and histopathologic evaluations. People who took part in this study had early-stage lesions that were thought to be signs of lymphocytic (LPP, FFA, CCCA), neutrophilic (TF, FD), or mixed (DC) PCA. In order to conduct pathway analysis and Real-time PCR analysis, a total of two 4-mm scalp biopsies were obtained from each patient. One biopsy was acquired from the afflicted region of the scalp, while the other was obtained from an unaffected region. To establish control conditions, we procured scalp biopsy specimens from a cohort of healthy individuals who were matched based on age and sex. The normal controls that were chosen did not display any indications of hair or skin abnormalities. All participants had symptoms of active disease, including erythema, discomfort, gradual alopecia, and indications of inflammation. The participants in this study were individuals who were 18 years of age or older, and they willingly offered their informed consent. A thorough assessment was undertaken, encompassing a review of the patient’s medical history, a comprehensive questionnaire pertaining to hair, and an inspection of the hair, scalp, and skin. The biopsies were conducted following the necessary ethical procedures, including receiving approval from the institutional human ethical committee and obtaining informed consent from all patients and volunteers involved. The tissue samples were stored at a temperature of −80°C until they were subjected to further processing. The biopsies were employed to extract total RNA, do pathway analysis, and perform real-time PCR.

### 2.2. Human Hair Follicle Outer Root Sheath Cells (HHORSCs)

Human Hair Follicle Outer Root Sheath Cells (HHFORS) were purchased from Science Cell Research, USA. The cells were seeded in 100mm plates at 0.6×10^6^ density. Experimental treatments included 25 µM 7-Dehydrocholesterol (7DHC) and 4 µM BM15766, with Ethanol and Dimethyl Sulfoxide (DMSO) as vehicle controls (<0.1% final concentration).

### 2.3. Animal study

The study involved doing experiments (IAEC 1-KU-12/21-ZOO-SRP (4)) on C57BL/6 mice that were seven weeks old. The mice were allocated into four groups, namely Control, vehicle, 7DHC, and BM15766, with each group comprising six mice. The depilation process was employed to achieve synchronisation of hair development by preserving hair follicles in the telogen phase. The scalp devoid of hair was subjected to topical application of a solution containing 25 millimolar (mM) 7DHC and 4 mM BM15766, which were dissolved in ethanol and dimethyl sulfoxide (DMSO), respectively, for a duration of 15 days.

### 2.4. Human hair follicles

The hair follicles (HFs) were sourced from authorized hair clinics, ensuring the ethical and regulated acquisition of these samples for our research. The hair follicle (HF) stands out among the various mini-organs within the human body due to its exceptional accessibility for experimental manipulation and ease of complete removal. Interestingly, even when removed from the body, the HF retains some essential in vivo characteristics within hair follicle organ culture (HFOC). Furthermore, the HF is unique among mammalian organs undergoing rhythmic and spontaneous cyclical transformations throughout the host’s lifespan. These transformations involve transitioning from a relatively dormant state (telogen) with minimal hair shaft formation and reduced HF size, to prolonged phases of robust and rapid growth.

Anagen VI, terminal hair follicles, have shown successful ex vivo growth for a duration of up to two weeks. To achieve this, the micro dissected hair follicles should be cultured in a completely defined, serum-free medium, Williams’ E medium. This medium was further enhanced by adding L-glutamine, hydrocortisone, insulin, penicillin, and streptomycin. The cultures were preserved at a constant temperature of 37°C in an atmosphere comprising 5% CO2 and 95% air (Paus & Cotsarelis 1999).After acclimatization, the hair follicles were grouped into four, control, vehicle, and two treated groups (25mM 7DHC and 4mM BM15766) and the compounds were treated continuously for four days.

### 2.5. RNA Isolation and pathway analysis

The total RNA from each biopsy was extracted using TRIzol reagent (BR Biochem Life Sciences, India) following the manufacturer’s instructions. The Ingenuity Pathways Analysis (IPA) software was used to discern the functional importance, cellular localization, and functions within the designated gene products’ diverse biological and metabolic processes.

### 2.6. Quantitative Real-Time RT-PCR

We performed RNA extraction following the TRIZOL RNA extraction protocol. SYBR Green-Labelled PCR primers for all targeted genes were purchased from G-Biosciences made in (St. Louis, MO, USA, Cat # 786-5062). Expressions of all targeted genes in PCA, normal scalp samples, HHFORS, mice, and HFs were quantified.

### 2.7. Immunofluorescence

Human Hair Follicle Outer Root Sheath Cells seeded in coverslip for culture. The cells were incubated for 24-48 hours and treated with 25mM 7 DHC, 4mM BM15766 for 24 hours. The cells then underwent washing, fixing, and permeabilization (Wu *et al*., 2014) and were incubated with targeted primary antibodies SOX 9, LGR5, and Wnt 5A, purchased from ImmunoTag (St. Louis, MO, USA, Cat# ITT02816). Subsequently, the cells were exposed to DAPI (1µg/ml) as a nuclear tagging agent.

### 2.8. BrdU Pulse-Chase experiment

Seven-week-old mice were selected for the BrdU pulse-chase experiment (BrdU kit, Invitrogen – Cat no. 8800-6599). The mice were divided into four groups for the experiment: control, vehicle (DMSO/ethanol), 7DHC, and BM15766. The removal of the superficial hair of the mice facilitated the synchronisation of their hair development cycle. The administration of BrdU commenced on the 21st day, which marked the initiation of the anagen cycle. This was followed by a chase period of 6 weeks, after which a second administration of BrdU was conducted on the 56th day. To administer an intraperitoneal injection, a solution containing a concentration of 1 mg of BrdU was generated by diluting it in 200 μL of sterile phosphate-buffered saline (PBS) at a concentration of 5 mg/mL. Subsequently, the mice were administered an intraperitoneal injection of 200 μL (1 mg) of the BrdU solution. The mice were subjected to careful monitoring during the course of the trial. Following the tracking phase, the mice were subjected to euthanasia, and subsequent tissue collection was conducted for the purpose of BrdU analysis.

### 2.9. TUNEL Assay

The tissues from treated mice and HFs were used for TUNEL apoptotic detection (In Situ Cell Death Detection Kit, TMR Red, Roche 12156792910) After fixation and antigen retrieval procedures, the tissue sections were subsequently exposed to the complete TUNEL reaction mixture and incubated in darkness for 60 minutes at 37°C. In the case of negative control tissues, only the labelling solution was applied, omitting the addition of terminal transferase. Conversely, the DNase I working solution was incorporated for positive control before proceeding with the labelling steps. After the designated incubation period, the counterstaining step was performed using DAPI at a 5 µg/ml concentration.

### 2.10. Immunostaining for hair follicle

All primary antibodies were obtained from ImmunoTag (St. Louis, MO USA). Cryosectioned hair follicles were permeabilized with 0.1% PBST and underwent antigen retrieval with a 10mM sodium citrate buffer at pH 6. Blocking was achieved using 5% goat serum. Incubation with primary antibodies was conducted overnight at 4 °C. Subsequently, the slides were washed, followed by incubation with secondary antibodies and counterstaining with DAPI.

### 2.11. Statistical Analysis

All experimental results were analysed by one-way analysis of variance (*p*<0.05) using SPSS software (Ver. 22.0). The data were reported as means with standard deviation, and a P-value below 0.05 was considered statistically significant.

## 3. Results

### 3.1. Down-regulated expression of stem cell marker genes (SOX9, LGR5, SHH, WNT 5A) in PCA

The data clearly demonstrated that the expression levels of the four target genes were consistently reduced across all subgroups of PCA. In Fig. 1A, it can be observed that the expression of SOX9 is significantly reduced in samples of Folliculitis Decalvans (DF), Dissecting Cellulitis (DC), Lichen Planopilaris (LPP), and Central Centrifugal Cicatricial Alopecia (CCCA) as compared to the control samples. The results of this study indicate a significant decrease in the expression of the LGR5 gene in the following conditions: TF, DF, DC, FFA, LPP, and CCCA, as shown in Figure 1B. The SHH and WNT 5A expression patterns are depicted in Figures 1C and 1D, respectively. The expression of SHH was found to be greatly increased in the samples of TF, whereas it was observed to be decreased in the impacted samples of DF, DC, FFA, LPP, and CCCA. Results are represented as mean ± SD (*P ≤ 0.05. **P ≤ 0.01). The affected samples show significant downregulation of the stem cell marker genes compared with the unaffected.

**Fig. 1.**
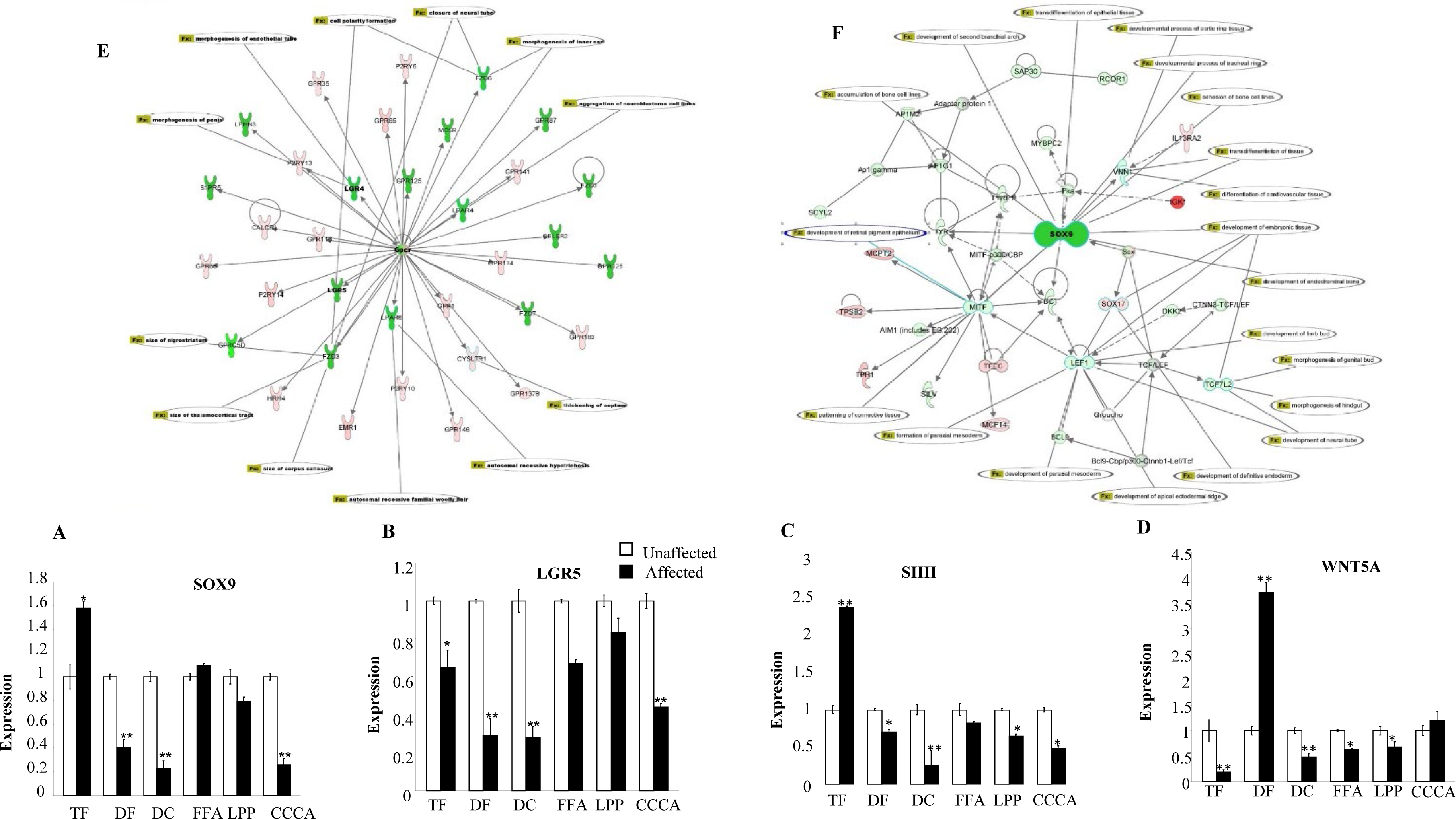
Downregulated expression of HFSC marker genes in PCA. Genes directly and indirectly connected with **(A)** LGR5 and **(B)** SOX9 are down-regulated. Nodes, genes; green nodes, genes down regulated in pathway; pink nodes, genes up regulated. Line between two nodes represents the corresponding protein products of the genes can physically interact (according to literature). The difference in intensity reflecting the degree of change in the expression of differentially expressed genes in our database. The difference in intensity reflecting the degree of change in the expression of differentially expressed genes in our database. Compared with the unaffected tissue, **(C)** SOX 9 gene expression was significantly downregulated in all PCA subtypes DF, DC, LPP, CCCA but not in TF and FFA. The un paired t-test used for statistical analysis. **(D)** Real time PCR validation of LGR5 gene expression in unaffected and affected skin in the PCA subtypes TF, DF, DC, FFA, LPP, CCCA (*p<0.05, **p<0.01). Compared with the unaffected tissue, LGR5 gene expression was significantly downregulated in all PCA samples. The un paired t-test used for statistical analysis. **(E)** Real time PCR validation of SHH gene expression in unaffected and affected skin in the PCA subtypes TF, DF, DC, FFA, LPP, CCCA (*p<0.05, **p<0.01). Compared with the unaffected tissue, SHH gene expression was significantly downregulated in all PCA subtypes DF, DC, FFA, LPP, CCCA but not in TF. The un paired t-test used for statistical analysis. **(F)** Real time PCR validation of WNT 5A gene expression in unaffected and affected skin in the PCA subtypes TF, DF, DC, FFA, LPP, CCCA (*p<0.05, **p<0.01). Compared with the unaffected tissue, WNT 5A gene expression was significantly downregulated in all PCA subtypes TF, DC, FFA, LPP, CCCA but not in DF. The one way ANOVA was used for the statistical analysis.

### 3.2. Pathway analysis of LGR5 and SOX9

The Gene interaction profile of the LGR5 pathway (Fig. 1E) included more than 30 genes and their significant functional regulations, such as morphogenesis, polarity, and neuronal regulations, etc. Usually, the hair follicle cycling is fuelled by stem cells. Genes directly and indirectly connected with SOX9 were down-regulated (Fig. 1F).

### 3.3. 7DHC or BM15766 inhibits the expression of SOX9, LGR5, WNT5A, and SHH in cultured human hair follicle outer root sheath cells (HHORS)

The experiment’s setup involved the administration of 7DHC and BM15766 treatments to HHORS cells, followed by subsequent analysis using real-time PCR. The results demonstrated a notable decrease in the expressions of SOX9, LGR5, WNT5A, and SHH in the treated samples (Fig. 2A–G) compared to the control group. The expression of LGR5 exhibited a significant decrease following treatment with both 7DHC and BM15766, as depicted in Figure 2B. Regarding the expression pattern of WNT5A (as depicted in Figure 2D), it was observed that its activity was downregulated upon treatment with 7DHC. However, no substantial downregulation was observed in the Sample treated with BM15766. The results were expressed as the mean ± SD(*P ≤ 0.05, **P ≤ 0.01).

**Fig. 2.**
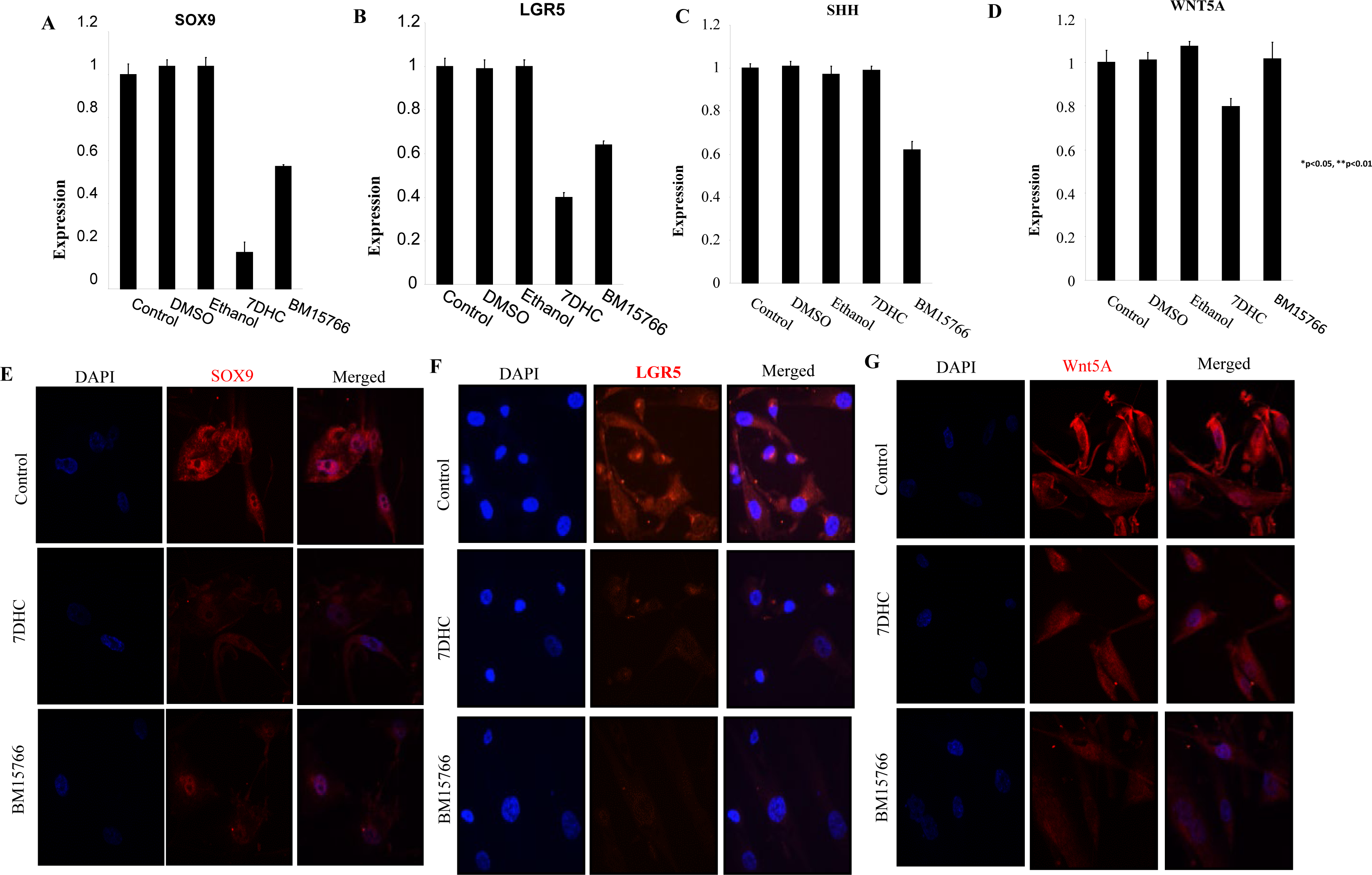
Inhibited cholesterol biosynthesis downregulates hair follicle stem cell marker genes in in vitro model system. The real-time PCR validation of **(A)** SOX9 **(B)** LGR5 **(C)** WNT 5A **(D)** SHH gene expression in 7-DHC and BM15766-treated samples, HFORSCs (*p<0.05, **p<0.01) is shown. Compared with the controls, treated samples show significant down-regulation in the expression of SOX9 and LGR5. The expression of WNT5A, down-regulated in 7DHC. SHH expressions are down-regulated in BM15766 treated Sample but not in 7-DHC. The one way ANOVA was used for the statistical analysis. Treatment with 7-DHC and BM15766 can reduce the expression of some or all of these genes. . Immunofluorescence analysis showed that **(E)**SOX9, **(F)** LGR5 and **(G)** Wnt 5A (Red) expressions are down-regulated compared with the control. Staining for the merge of SOX9, LGR5 and WNT 5A with DAPI respectively.

The RT-PCR results were further corroborated through protein expression. The immunofluorescence analysis revealed that, in comparison to the untreated samples, the cells expressing with SOX9, LGR5, and WNT5A were noticeably reduced in the treated samples. Specifically, Fig. 2E depicts the diminished expression of SOX9 after treatment with 7DHC and BM15766. In the case of LGR5 (Fig. 2F), the treated images demonstrate reduced expressions compared to the control. Similar reductions were observed in the expressions of WNT5A proteins (Fig. 2G), as seen with SOX9 and LGR5.

### 3.4. 7DHC and BM15766 halt hair regrowth by altering cholesterol biosynthesis

After administering a 15-day treatment of (Fig. 3A)7DHC and (Fig. 3B) BM15766 on the mouse scalp, a comprehensive absence of hair regrowth was detected, suggesting the robust inhibitory properties of both compounds. Based on the observed outcome, it was postulated that the diminished hair regeneration could potentially be linked to modified expression patterns of hair follicle stem cell marker genes in the treated samples. To substantiate this hypothesis, we performed a reverse transcription polymerase chain reaction (RT-PCR) study on scalp tissues from mice, specifically targeting prominent stem cell markers such as SOX9, LGR5, SHH, and WNT5A.

**Fig. 3.**
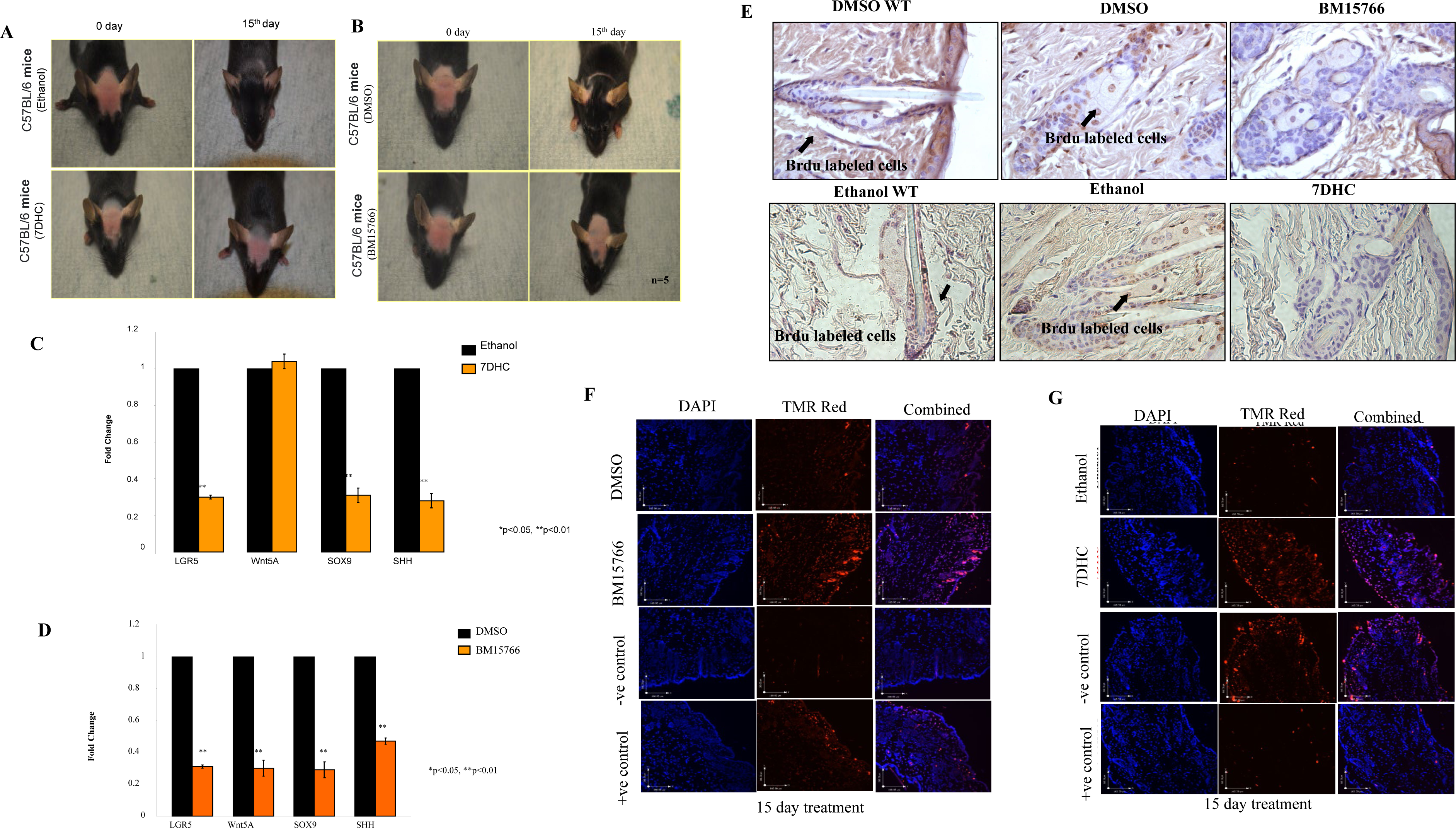
7DHC and BM15766 treated mice (C57BL/6; n = 5) failed to regrow the hairs. Treatment with topical **(A)**7DHC and **(B)** BM15766 resulted in a reduction in hair growth when compared to mice treated with Ethanol/ DMSO. However, while hair growth was successfully restored in the mice treated with the vehicle, it did not recover in the treated group. Real-time PCR validation of LGR5, WNT 5A, SOX9 and SHH gene expression in mouse skin treated with **(C)** 7DHC and **(D)** BM15766 compared with vehicle-treated (Ethanol/ DMSO) controls (n = 5; *p,0.05, **p,0.01). Treatment group can downregulate the expression of some or all of these genes. **(E)** Labeling with BrdU during the pulse phase and the subsequent chase period, the experiment did not yield conclusive evidence of stem cell populations retaining the BrdU label in 7DHC and BM15766 treated mice. In contrast the control (Ethanol/ DMSO) tissues the experiment showed the presence of BrdU labeling. TMR red imaging revealed a higher number of apoptotic or DNA damaged (red color) cells in the **(F)** BM15766 and **(G)** 7DHC treated samples compared to the +ve and vehicle (Ethanol/ DMSO) control group. Tissue counterstained with DAPI(nucleus)

The RT-PCR study confirmed our initial hypotheses, as evidenced by the decreased expression of stem cell marker genes in the (Fig. 3C)7DHC and (Fig. 3D) BM15766 treated samples compared to the vehicle control. This observation provides more evidence to support the hypothesis that disruptions in the process of cholesterol production could potentially contribute to the compromised ability of hair to regrow and the detrimental effects on hair follicle stem cells. The potential contribution of sterol intermediate buildup resulting from the interruption of cholesterol production pathways to the observed damage in stem cells is worth considering.

### 3.5. *In vivo* 7DHC and BM15766 Treated Samples Exhibits Stem cell Exhaust

To identify the fate of stem cells, we conducted a BrdU pulse-chase experiment, which aids in locating label-retaining cells in the *in vivo* (Fig. 3E) system. During the BrdU pulse phase and the subsequent chase period, the experiment failed to provide conclusive evidence of stem cell populations retaining the BrdU label in BM15766 and 7DHC treated mice. However, in stark contrast, the control tissues treated with DMSO/ethanol exhibited the presence of BrdU labelling. The absence of BrdU-labelled cells in the BM15766 and 7DHC treated mice suggests these compounds might adversely affect stem cell function and differentiation. A plausible hypothesis posits that the suppression of cholesterol production and the buildup of sterol intermediates have a detrimental impact on the functionality of stem cells. Cholesterol and its derivatives are key components in many cellular functions, encompassing the establishment and maintenance of cell membrane integrity, intercellular communication, and the regulation of gene expression. Impairment of these pathways may result in compromised maintenance and proliferation of stem cells.

### 3.6. Increasing Proportion of "TUNEL" Positive Cells in *In vivo* 7DHC and BM5766 treated Sample

The disappearance of label-retaining cells in the presence of 7DHC and BM15766 raises intriguing questions about the fate of stem cells and the possibility of apoptosis. The presence of TUNEL-positive cells would provide strong evidence of apoptotic events in the stem cell population. The TMR red imaging demonstrated a notably elevated count of apoptotic or DNA-damaged cells (indicated by the red colour) in the BM15766 (Fig. 3F) and 7DHC (Fig. 3G) treated samples when compared to both the positive control group and the vehicle (ethanol) control group. The tissue was counterstained with DAPI to visualize cellular structures, highlighting the nuclei accurately.

### 3.7. Effect of inhibited cholesterol biosynthesis and intermediates on human hair follicle organoid culture

The hair follicles were exposed to 7DHC and BM15766 for four days and evaluated the stem cell changes (Fig. 4A). No apparent alterations were observed in the hair follicles of the control group during the investigation. Nevertheless, a notable disparity was detected among the group that received treatment, as the length of the follicles exhibited a substantial decrease. The formerly orderly and condensed arrangement exhibits signs of deformation, indicating possible impairment of the stem cell microenvironment. Additionally, there are indications of degradation in the hair matrix, the crucial area responsible for initiating hair fibres. The presence of structural modifications, disrupted cellular arrangement, and degradation of the hair matrix collectively suggest potential disturbances in the functionality of stem cells and the process of hair regeneration.

**Fig. 4.**
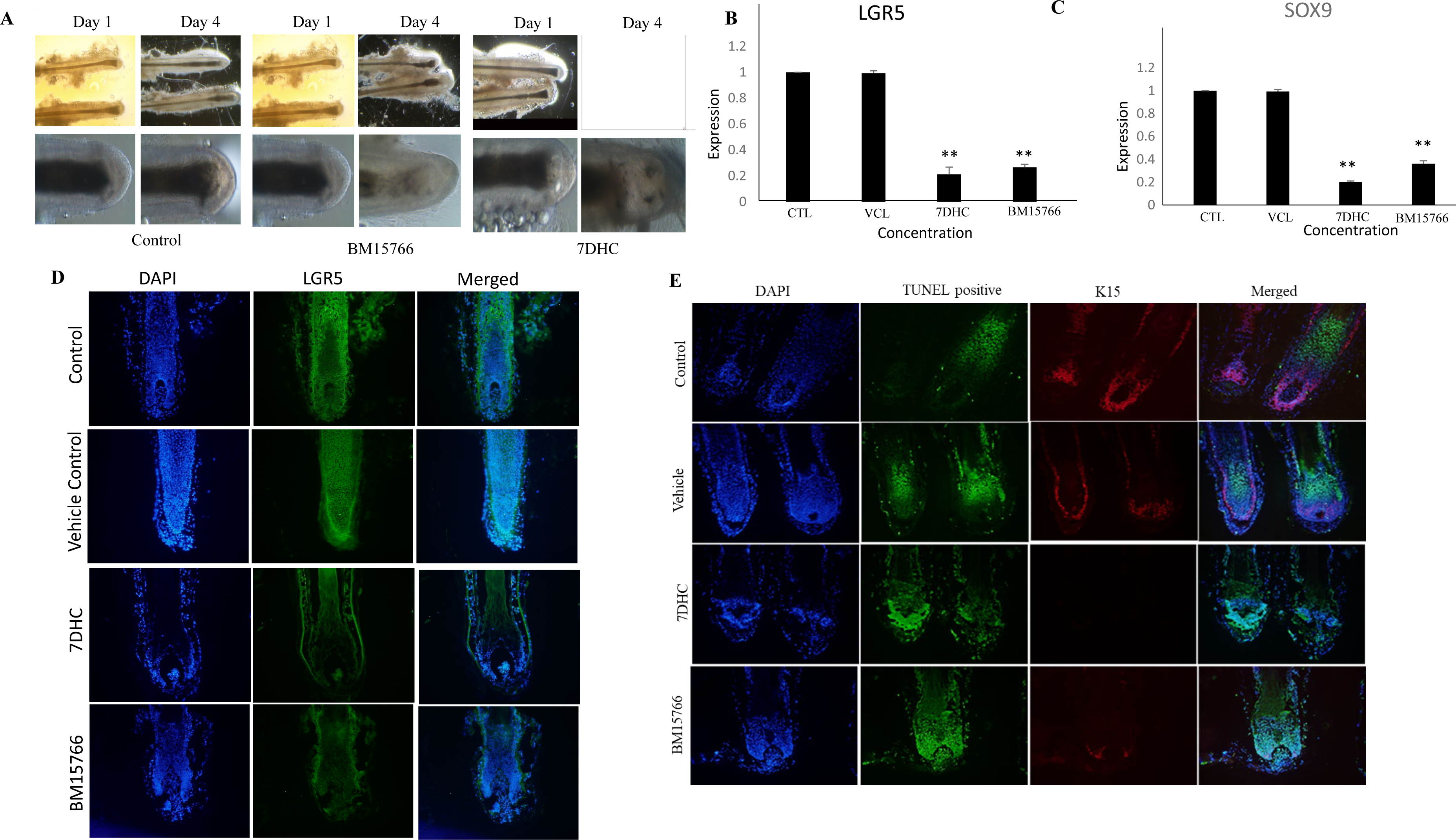
Inhibitory effect of 7DHC and BM15766 in HF organoid culture. **(A)**The diagram provides a visual representation depicting the detrimental effects of BM15766 and 7DHC on the hair follicle bulge. The hair follicle bulge, a critical region housing stem cells. The detrimental impact on the bulge region is indicated by structural alterations, disruption of cellular organization, deterioration of hair matrix and the hair fiber pushed upwards within the follicle. The real-time PCR validation of **(B)** LGR5 and **(C)** SOX9 gene expression in 7-DHC and BM15766-treated samples, HF (*p<0.05, **p<0.01) is shown. Compared with the controls, treated samples show significant down-regulation in the expression of SOX9 and LGR5 The one way ANOVA was used for the statistical analysis. **(D)** Immunofluorescence imaging of the LGR5 (Alexa fluor 488) in HFs with DAPI counter staining. Compared with the control, treated samples shows reduced expression of LGR5. **(E)** Immunofluorescence imaging of the expression of K15(Alexa fluor 594) and TUNEL positive cells, of HFs treated with 7DHC and BM15766. The HFs are counter stained with DAPI and finally merged. The treatment group showed higher apoptotic cells than the control group. Similarly the expression of K15 was drastically reduced when the HFs are exposed to 7DHC and BM15766.

**Fig. 5.**
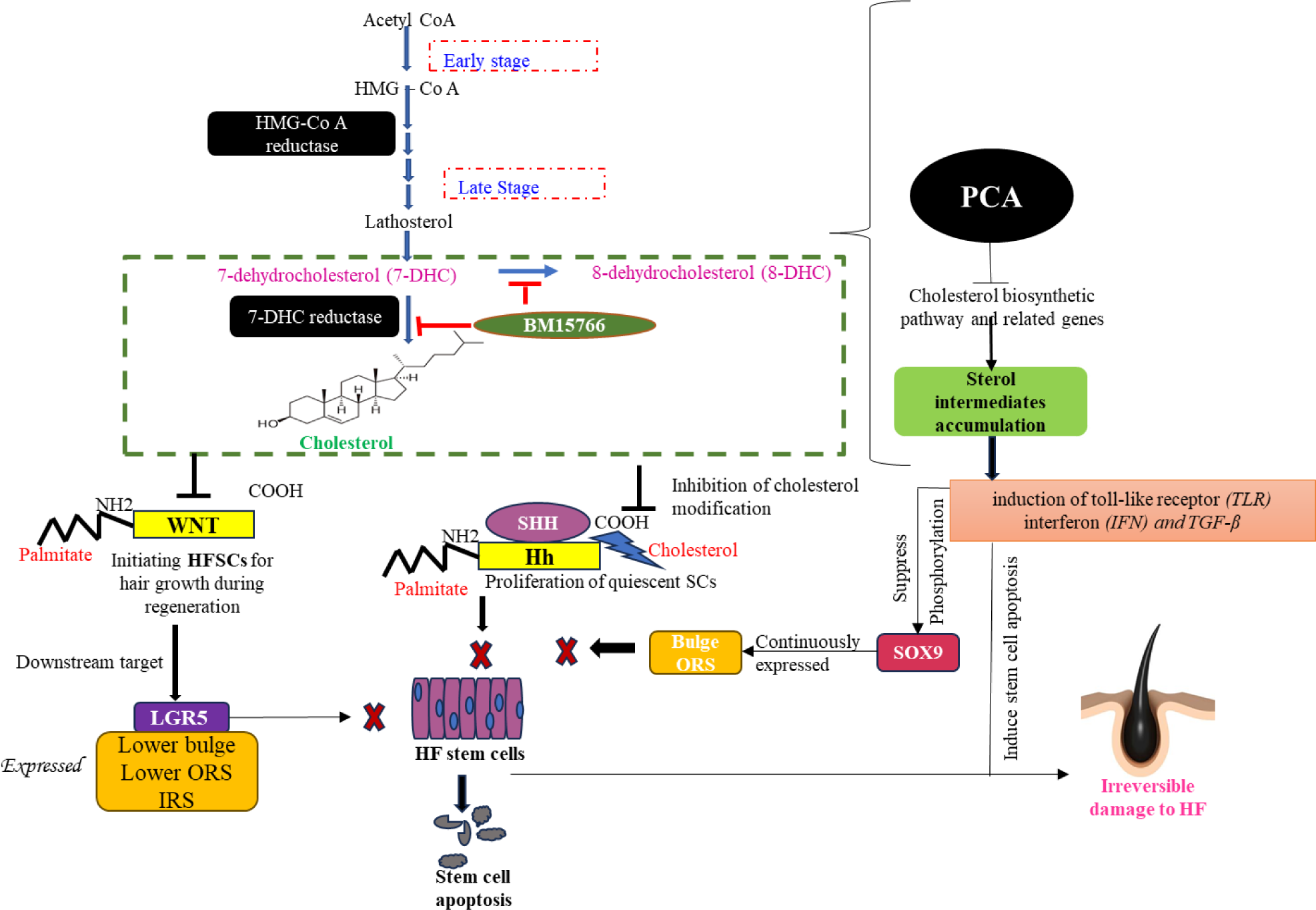
In individuals affected by PCA, there is a significant downregulation of genes associated with the cholesterol biosynthetic pathway. This downregulation results in the accumulation of sterol intermediates, which triggers inflammatory responses within hair follicle (HF) stem cells, ultimately leading to the apoptosis of these stem cells. The inflammatory reactions involve various elements, such as Toll-Like Receptors (TLR), Interferons (IFN), and Transforming Growth Factor-beta (TGF-β). These inflammatory factors collectively have a suppressive effect on the phosphorylation of SOX9, a crucial protein in stem cell proliferation, particularly in the bulge of the Outer Root Sheath (ORS). This suppression further contributes to damage to the stem cells. To understand the mechanistic aspects of this process, we introduce two key compounds, 7-Dehydrocholesterol (7DHC) and BM15766, which artificially alter the cholesterol biosynthetic pathway. These compounds effectively mimic the conditions experienced by PCA-affected individuals.In this altered state of the cholesterol biosynthetic pathway, we find a significant connection with critical stem cell markers, specifically Hh, WNT, LGR5. The N-terminus of Hh is modified by adding palmitate to a conserved cysteine residue, while the C-terminus is modified with cholesterol. Similarly, WNT proteins undergo palmitoylation at their first conserved cysteine. These lipid modifications are crucial for the precise localization of both WNT and Hh proteins and further effective functioning in the development and cycling of hair follicles.In stem cell regeneration, Hh plays a pivotal role in promoting the proliferation of quiescent stem cells, ultimately leading stem cell regeneration. Similarly, WNT signaling is instrumental in regenerating Hair Follicle Stem Cells (HFSCs) during hair growth.The absence of cholesterol modification in Hh and WNT due to the altered cholesterol biosynthetic pathway disrupts stem cell proliferation. Moreover, this disruption affects downstream targets of the WNT pathway, such as LGR5, leading to the failure of its expression in the lower bulge, ORS, and IRS regions. Coupled with the presence of inflammatory cells, this disruption can culminate in the apoptosis of stem cells, ultimately contributing to the pathology observed in PCA-affected individuals.

### 3.8. Human hair follicle stem cell markers were downregulated in *ex vivo* culture

The inhibited cholesterol biosynthetic pathway has demonstrated its ability to impact the functioning of hair follicle stem cells. The obvious downregulation of stem cell markers serves as evidence that this claim has support from both *in vitro* and *in vivo* model systems. The results were highly promising and further confirmed the negative regulation the inhibited cholesterol biosynthetic pathway imposed on hair follicle stem cell marker genes, including (Fig. 4B) LGR5, (Fig. 4C) SOX9. Notably, these marker gene expressions were significantly downregulated, as evidenced by the results presented as mean ± SD (*P ≤ 0.05, **P ≤ 0.01).

### 3.9. Unlocked the Secrets of Hair Follicles: 7DHC and BM15766 Treatments Revealed Downregulated LGR5 and Apoptosis of K15

Our *in vitro* and *ex vivo* experiments showed that LGR5 expression decreased significantly when cholesterol biosynthesis was stopped. Remarkably, we were able to replicate these similar results during *ex vivo* organ culture, strengthening the robustness of our findings. Consequently, the evidence was unequivocal that follicles treated with 7DHC and BM15766 experienced a pronounced deterioration in LGR5 (Fig. 4D) stem cell population, ultimately exerting a substantial influence on the overall hair follicle growth and cycling process. In addition to its presence in the bulge area of hair follicle cells, K15 is a crucial indicator of stem cell populations. Our results revealed a significant downregulation of K15 (Fig. 4E) expression in the treated samples, along with the identification of TUNEL TMR red expressions, indicating the presence of apoptotic cells.

## 4. Discussion

The study reveals how cholesterologenic changes and subsequent sterol intermediate accumulation might be responsible for the irreversible damage to hair follicles in individuals affected by PCA. Earlier studies have suggested that the persistent destruction of scalp hair follicles in PCA-affected individuals may be attributed to the depletion of hair follicle stem cells (Cotsarelis and Miller, 2001). Our current study provides robust support for these findings, as our data reveals consistent downregulation of components in the network analysis of signalling pathways (SOX9 & LGR5). Moreover, specific stem cell marker genes, including SOX9, LGR5, SHH, and WNT5A, were significantly downregulated in the scalp of PCA patients, further corroborating our results. Karnik *et al*., 2009 have extensively studied the relationship between PCA and cholesterologenic changes. Their findings suggest that the targeted deletion of PPARγ (a transcription factor involved in regulating lipid homeostasis) in follicular stem cells leads to scarring alopecia. PPARγ significantly influences lipid biosynthesis and inflammatory responses (Karnik et al., 2009). Notably, PCA patients exhibit a significant decrease in the expression of this transcription factor, indicating its role in lipid biosynthesis and hair follicle cell functioning. Consequently, cholesterologenic changes are deemed to play a crucial role in the inflammatory responses observed in individuals affected by PCA (Panicker et al., 2012). Based on this data, regulating cholesterol biosynthetic genes and hair follicle stem cell functioning are considered hallmark features of PCA. Hair follicle stem cells are paramount in PCA pathology (Harries et al.,2013). When these stem cells are affected for any reason, hair follicles are destroyed, rendering them incapable of regrowth. Subsequently, the affected hair follicles enter the catagen stage, culminating in the formation of a fibrotic tract and scarring alopecia (Whiting & Olsen., 2008).

Prior research has consistently identified LGR5 as a reliable marker for hair follicle stem cells (HFSC) in the bulge region (Yang et al., 2014). LGR5 and its associated signalling molecules play a crucial role in the maintenance and development of hair follicles. Proper LGR5 pathway activation is essential for the continuous renewal of hair follicles and hair regrowth after shedding. However, in individuals affected by PCA, the LGR5 pathway becomes downregulated or has reduced expression. This may result in the depletion or destruction of these stem cells, leading to a reduced ability to regenerate hair. Building upon this previous knowledge, we sought to delve deeper into understanding how changes in cholesterol homeostasis might influence hair follicle stem cell dynamics in PCA. In our in vitro analysis we observed significant downregulation of hair follicle stem cell markers upon treatment with 7DHC and BM15766. Recent studies have revealed that hair follicles have the capacity for intrafollicular de novo biosynthesis of cholesterol, with a key enzyme, 24-dehydrocholesterol reductase (DHCR24), being highly expressed in these follicles. 24-dehydrocholesterol reductase is crucial in converting desmosterol to cholesterol, contributing to the cholesterol levels necessary for proper hair follicle function (Brannan et al., 1975; Mizra et al., 2009).

So, the altered cholesterol biosynthetic pathway using 7DHC and BM15766 also satisfied the previous study’s report that the inhibited cholesterol biosynthetic pathway induces the accumulation of sterol intermediates and leads to inflammatory infiltration and further hair loss (Panicker *et al*., 2012). While the impact of lipids on hair follicle (HF) biology is well recognized, the precise functions influenced by cholesterol still need to be clarified. Cholesterol’s connection with the lipid-rich sebaceous gland in HF suggests its potential significance. Although some associations have been identified between sterol levels and specific hair disorders, the exact role of cholesterol in hair health requires further investigation. Additionally, drug therapies that modulate lipids have been reported to affect hair loss and growth, indicating the complex interplay between lipids and hair physiology (DeStefano et al., 2014; Evers et al., 2010).

The mice receiving 7 DHC and BM15766 treatments showed clear signs of epidermal thickening, fibrosis, and a failure to regrow hair. Lipids and their role in hair follicle (HF) biology are crucial, and cholesterol, in particular, has long been suspected of significantly influencing hair growth. We now understand that disruptions in cholesterol balance are intricately tied to conditions like PCA, where hair follicles are permanently damaged, and congenital hypertrichosis, characterized by excessive hair growth due to mutations in cholesterol transporters. Moreover, there’s a fascinating connection between abnormal lipid levels in the body, known as dyslipidemia, and androgenic alopecia, a common form of hair loss linked to hormones (Palmer et al., 2020). Cholesterol plays a crucial role in the physiology of the skin by serving as a fundamental component in maintaining the integrity of the epidermal barrier (Wertz, P. W. 2000) and serving as a precursor for the synthesis of steroid hormones (Slominski, et al., 2013). Cholesterol modifications play a crucial role in facilitating signal transduction within the Wnt-β-catenin and hedgehog pathways (Incardona and Eaton 2000), which are essential components in regulating the human hair follicle (HF) cycling process (Lee and Tumbar 2012).

Additionally, the observed downregulation of stem cell markers in the treated samples provides further insight into their fate. For example, the N-terminus of Hedgehog (Hh) is altered by adding a fatty acid palmitate to a conserved cysteine residue, while the C-terminus is modified with cholesterol. In the case of WNTs, they undergo palmitoylation on the first conserved cysteine (C77), which is present in all WNT molecules. Genetic evidence indicates that these lipid modifications are essential for localizing WNT and Hh proteins to cell membranes, a prerequisite for their effective function in hair follicle development and cycling (Nusse., 2003). An experiment involving a targeted mutation in the mouse’s endogenous Hh gene, specifically the sonic hedgehog gene, revealed that when cholesterol modification is disrupted, it leads to a reduction in the effective range of Hh activity, potentially even resulting in a complete loss of Hh function (Lewis et al., 2001). Loss of PPARγ (Wan et al., 2007; Karnik et al., 2009) or the accumulation of sterol precursors (Evers et al., 2010) can lead to disruptions in fatty acid metabolism and cholesterol biosynthesis. These disruptions can hinder the proper lipid modification of Wnt and Hh proteins, potentially causing interference with hair follicle development and cycling. Through the TUNEL apoptotic assay, it became evident that there was an increased count of apoptotic cells within the treated samples. This finding aligns with the observed decline in stem cells during the label retention pulse-chase experiment.

Our study elucidated the functional characteristics of stem cells at the molecular level, revealing a significant observation regarding the decreased expression of crucial markers associated with hair follicle stem cells. Specifically, the markers SOX9, SHH, LGR5, and WNT 5A were found to be downregulated in hair follicles treated to 7DHC and BM15766. The downregulation described may play a significant role in elucidating the mechanism behind the irreversible degeneration of hair follicles reported in persons affected by PCA. When the expression of these markers is dramatically reduced, it seems to impede the intrinsic capacity of the hair follicle to undergo repair and regeneration. Consequently, the inflammatory process in primary cicatricial alopecia (PCA) causes irreversible damage to the hair follicles, resulting in permanent hair loss that cannot be restored through natural regeneration mechanisms. Comprehending these molecular pathways is of utmost importance in devising tailored interventions and therapies to address the fundamental aetiology of hair loss in individuals afflicted by PCA. This discovery presents opportunities for further investigation and therapeutic approaches focused on reinstating the normal expression of these stem cell markers which may facilitate the regeneration of hair follicles and promote hair growth.

Our research proves that the cholesterol biosynthesis pathway plays a significant role in maintaining hair follicles (HFs) and promoting the proper functioning of the stem cells residing inside them. Therefore, the potential for therapeutic intervention in the fields of alopecia and regenerative medicine lies in establishing a harmonious interplay between these two systems. The requirement stems from the recognition that the preservation and resilience of hair follicle stem cells (HFSCs) play a crucial role in sustaining optimal hair growth and mitigating disorders like alopecia. The primary objective of this study was to improve the survival of hair follicle stem cells (HFSCs), intending to address alopecia and advance our comprehension of tissue regeneration and the complex mechanisms that govern hair biology.

These endeavours have the potential to establish new approaches and strategies for enhancing hair health and aesthetic satisfaction in a sustainable manner.

## ACKNOWLEDGMENT

This research was funded by Kerala Biotechnology Commission, YIPB program, KSCSTE; HRD scheme, Dept. of Health Research-Start up Grant, Govt. of India, and the University of Kerala. We would also like to thank the technical staff at Advanced Centre for Regenerative Medicine and Stem cell in Cutaneous Research (AcREM-Stem) who generously contributed to this research.

